# “OmniBio: An Easy-to-Use Web App for Kinetic Growth Parameter Calculation from Microplate Reader Data”

**DOI:** 10.1101/2025.09.05.674595

**Authors:** Sebastian Dehnhardt-Amengual, Catalina Ardiles, Ignacio Guarda, Luis F. Larrondo, Wladimir Mardones

## Abstract

OmniBio is a user-friendly web application designed to streamline the analysis of microbial growth curves from microplate reader data, eliminating the need for coding skills or cumbersome file preprocessing. Built in R, using the gcplyr package and Shiny framework, OmniBio supports data outputs from both Gen5 and iControl software, handling multiple sheets within a single file. Users can upload raw data along with experimental metadata to the platform, compute kinetic parameters such as maximum growth rate, lag time, ODmax, and area under the curve, and promptly visualize the results in real-time. The app produces plots for each parameter and a global heatmap analysis for comparing microbial performance across strains and culture media. Importantly, all the primary and processed data can be readily downloaded in a summarized Excel report. In contrast to existing tools, OmniBio optimizes the analysis process by automating repetitive computations and offering an intuitive, step-by-step workflow that does not require bioinformatics training. OmniBio is freely accessible at https://sdehnhardt.shinyapps.io/OmniBio/, and can be modified from open-source code on GitHub (https://github.com/sidehnhardt/OmniBio). The tool enables researchers to conduct high-throughput kinetic analysis in an efficient and user-friendly manner, optimizing both time and resource utilization.

## Introduction

Bacterial growth curve analysis has been a fundamental strategy for understanding the physiological characteristics of microorganisms through the determination of their kinetic parameters (Fernández-Martínez et al., 2024). Typically, for this purpose, different kinetic parameters are compared across various culture media to determine phenotypic characteristics using a wide range of available software (e.g., GrowthRates or Microbial Lag Calculator) and R packages (e.g., GrowthCurver) (Hall et al., 2014; Sprouffske & Wagner, 2016). Nevertheless, the use of R packages or other bioinformatics tools requires some skillsets or training, which complicates their widespread use by all lab members. With the advent of robotics in laboratory procedures, the need to process extensive data generated by this type of equipment has arisen, normally compromising significant analysis time. In this context, bioinformatics pipelines represent a viable alternative for automating repetitive tasks. Considering this, our goal is to use the gcplyr R package (Blazanin, 2024), adapting it to handle multiple microplate readings of Optical Density in various culture media. Although the package already simplifies these tasks, we surmise that a user interface (UI) could provide a more efficient and real-time way to run the code in the backend while visualizing the results of the calculations. With this in mind, we developed OmniBio, which can read and analyze data from Gen5 and iControl software of any microplate reader, handling several sheets within a single Excel file. In this platform, users can visualize the results of calculations in real-time and evaluate the growth kinetics in each well of a microplate. Also available is a tab that stores the results for fast comparisons between culture media and a tab with a Heatmap analysis that summarizes each of the parameters across experimental sets, permitting global comparative analyses of microbial behaviors. Additionally, all the calculations performed by OmniBio can be downloaded in a summarized Excel file. This tool is freely available at https://sdehnhardt.shinyapps.io/OmniBio/.

## Workflow and features

One of the advantages of OmniBio is that, as hinted earlier, it does not require previous modification of the plate reader data output file. All calculations to determine the kinetic parameters are based on the code provided by the R package gcplyr (Blazanin, 2024). First, the user must select the type of input file to be uploaded: Gen5 or iControl. This option activates one of these two coding paths. For iControl data sheets, the user must add the number of the Label where the optical density measurements are located. This allows OmniBio to identify the position of data across the sheet (Figure 1).

**Figure 1.**
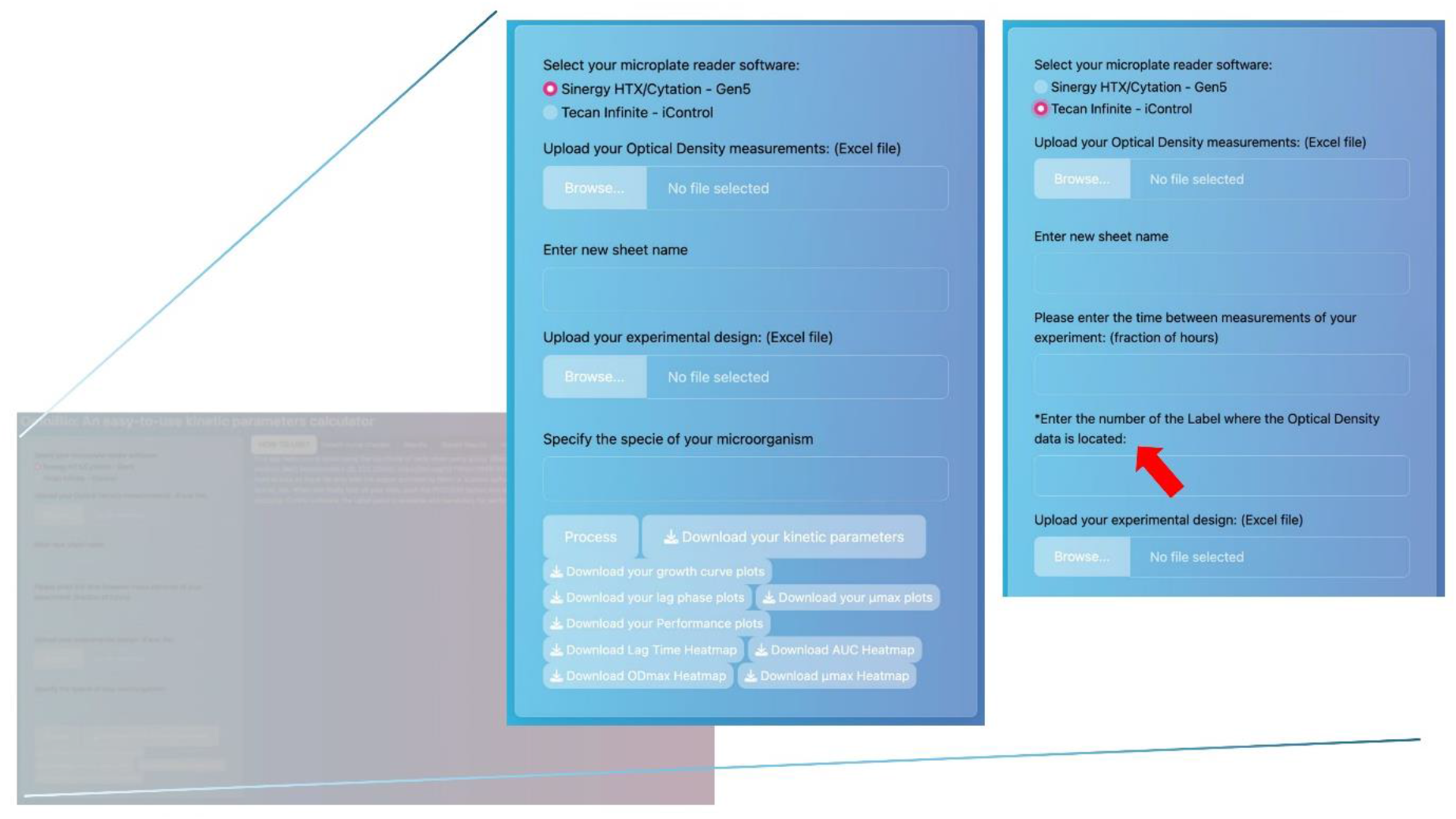
Snapshots of the two different coding paths for Gen5 or iControl analysis. The red arrow indicates the principal difference between the two types of output data software. Note that this option appears only when the Tecan Infinite-iControl option is selected in the first step.

Independent of the file type, a browsing list shows the available data sheets, allowing the user to perform multiple calculations of growth curve parameters using the same workbook. Note that the sheet name displayed corresponds to the one specified in the workbook. After selecting the sheet to be calculated, the user is required to enter a new name for the sheet, allowing the program to add it to the results file (it is allowed to enter letters and numbers in this box). The purpose of this new name is to serve as an identifier for the calculation results and to assign a clever reminder of each sheet analysis. This is useful if you are working with multiple culture media to understand the change in kinetic parameters for a particular microorganism. OmniBio next requires inputting the time between each measurement in a fraction of hours format (i.e., 30 minutes = 0.5 hours).

Furthermore, it is necessary to upload another file corresponding to the “metadata of the experiment”, which is independent of the well plate used for the experiment. This metadata serves as an identifier for the different samples (i.e., strains, isolates, media composition/condition) across the plate, allowing the program to assign a name to each well and perform further calculations for the kinetic parameters. Finally, the platform requires adding the name of the species of interest and selecting the “Process” button. It does not matter which name you specify in species, since this only allows you to add the name to the final Excel results.

After entering all the above requirements, the web-based program starts to perform its calculations. Depending on the file size and the number of analyzed wells, processing takes a few seconds to nearly one minute to complete. To verify that the calculations were performed correctly, please navigate to the “Growth curve checker” tab and expect to see the plotted curves. Another way to check if the processing is finished is to see if the download buttons are selectable.

OmniBio has various tabs to visualize the calculations and the values of the different kinetic parameters. Each of these tabs provides a brief description of the parameter, including its fundamentals. Note that the last three tabs (Area Under the Curve, All Parameters, and Stored Results) do not have any plot associated with them. To get the post-processing results in Excel, the user must select “Download Kinetic Parameters”. OmniBio can store the results of each sheet’s calculations, allowing the user to review and compare previous results with those from other sheets (e.g., comparing culture media).

One of the functional analyses for high-throughput data analysis is a global comparison across strains and media cultures. For this, the heatmap provides a comprehensive alternative for intuitive data analysis (Cremonini, 2024). The OmniBio platform can analyze the calculated data and generate a heatmap analysis for all kinetic parameters calculated. For this, the code compiles all the data generated across multiple sheets in a file, saves it in a temporary object, and scales the data to visualize it in a heatmap with a dendrogram for columns and rows, using the R base function “heatmap” (Figure 2). Indeed, in the provided example (File S1, S2 and S3), one can easily compare 16 yeast strains across 5 growth conditions, such as different carbon sources and osmotic stresses. The output figure permits comparing key growth parameters such as ODmax, lag time, µmax, and AUC. The heatmaps of the example data show that the control strain exhibits marked differences compared with the other strains, as evidenced by its separation into a distinct cluster (Figure 2). Similarly, maltose emerges as a condition that induces differential behavior compared with the other culture conditions tested.

**Figure 2.**
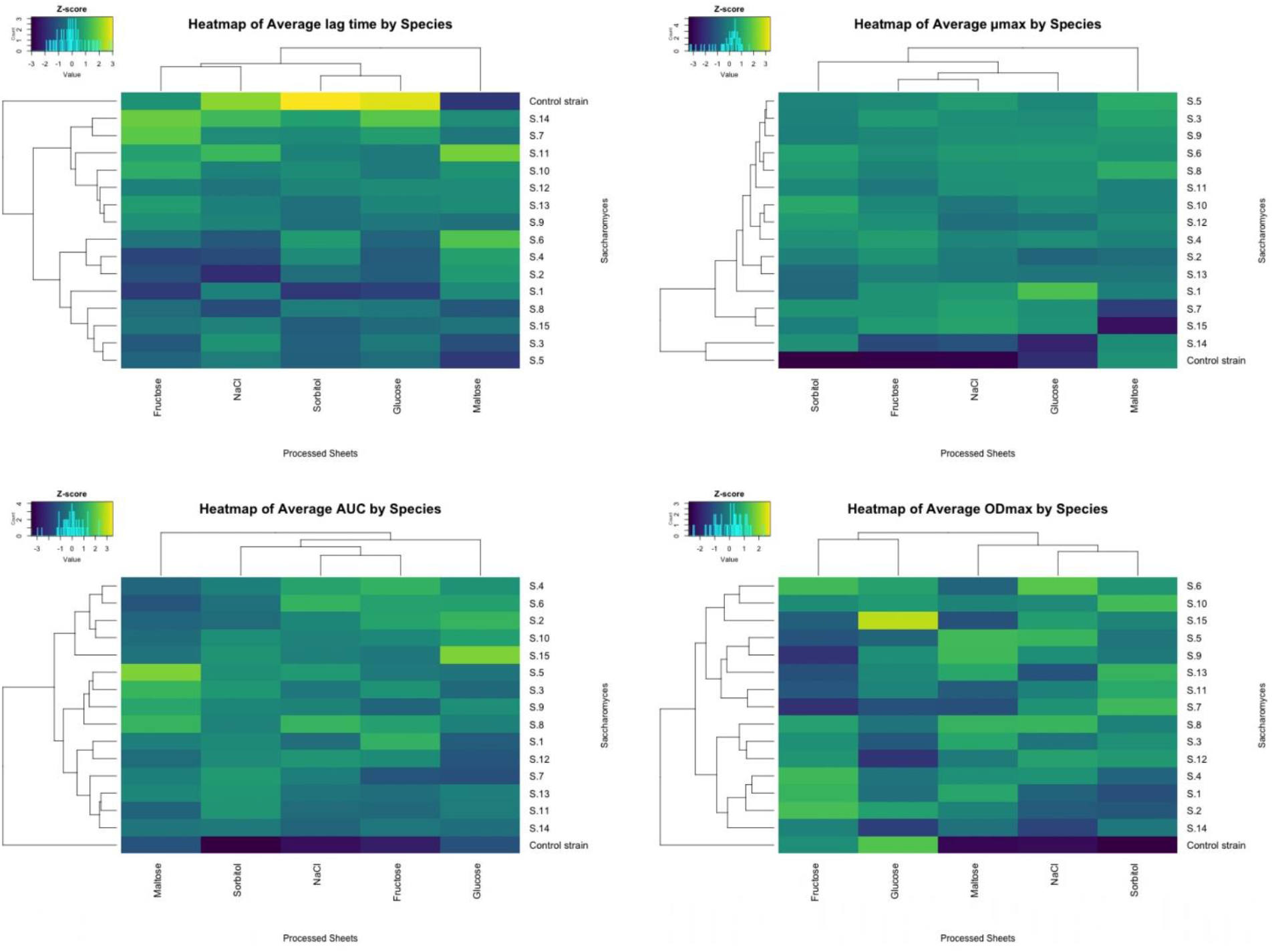
Heatmaps rendered by OmniBio with a multiple-sheet output file. In the Figure are shown four heatmaps, each corresponding to one of the kinetic parameters. The brighter color shows the higher Z-score value.

For more detailed instructions on how to use OmniBio, go to the supplemental material and watch the example video (Video S1). This material also includes sample files, including one Gen5 and iControl output, with the respective metadata examples.

OmniBio was developed entirely in the R language using RStudio and Shiny App programming logic. Furthermore, the complete code was published on the shinyapps.io server, accessible through the link: https://sdehnhardt.shinyapps.io/OmniBio/. The open-source code is fully available on GitHub (https://github.com/sidehnhardt/OmniBio).

## Concluding remarks and limitations

OmniBio is faster and user-friendly for analyzing high-throughput data retrieved from two of the most widely used microplate reader software platforms (Gen5 and iControl). Furthermore, the app can handle the data workbook exactly as it is exported from the software, avoiding any data pre-processing or modifications that could lead to undesired errors. However, OmniBio is currently unable to blank samples across the OD measurements, which will be implemented in future app updates. Although the software was originally developed for microbial growth curve analysis, it can also be applied to other types of response curves, such as luminescence signals from reporter assays, using the AUC parameter as a measure of induction. We are willing to perform code modifications that include calculating diauxic shifts and compiling all the results obtained into one Excel file. Finally, due to the open-source nature of OmniBio, customization and improvement of the functionality of the software could be implemented

## Acknowledgements and Funding

This work was supported by the National Association of Research and Development (ANID) with the FONDECYT Iniciacion Grant N° 11240430, FONDECYT Grant N° 1251234, and Programa Iniciativa Científica Milenio - ICN17_022. We are also grateful for the use of Cytation 3 (ANID/FONDEQUIP EQM130158) and BioStack4 equipment, as well as the Molecular Genetics Laboratory of the Santiago

University of Chile for supplying experimental data coming from Tecan Infinite plate reader and iControl software.

## Notes

### Competing Interest Statement

The authors have declared no competing interest.

## References

Blazanin, M. (2024). gcplyr: An R package for microbial growth curve data analysis. BMC Bioinformatics, 25(1), 232. 10.1186/s12859-024-05817-3

Cremonini, M. (2024). Heatmaps. In Data Visualization in R and Python (1st ed., pp. 157–163).

Fernández-Martínez, L. T., Javelle, A., & Hoskisson, P. A. (2024). Microbial Primer: Bacterial growth kinetics. Microbiology, 170(2), 001428. 10.1099/mic.0.001428

Hall, B. G., Acar, H., Nandipati, A., & Barlow, M. (2014). Growth rates made easy. Molecular Biology and Evolution, 31(1), 232–238. 10.1093/molbev/mst187

Smug, B. J., Opalek, M., Necki, M., & Wloch-Salamon, D. (2024). Microbial lag calculator: A shiny-based application and an R package for calculating the duration of microbial lag phase. Methods in Ecology and Evolution, 15(2), 301–307. 10.1111/2041-210X.14269

Sprouffske, K., & Wagner, A. (2016). Growthcurver: An R package for obtaining interpretable metrics from microbial growth curves. BMC Bioinformatics, 17(1), 172. 10.1186/s12859-016-1016-7

